# D-mannose ameliorates age-associated cellular senescence in the bladder urothelium and NLRP3/Gasdermin/IL-1β -driven pyroptotic epithelial cell shedding

**DOI:** 10.1101/2022.09.06.506836

**Authors:** Chetanchandra S. Joshi, Arnold M. Salazar, Caihong Wang, Marianne M Ligon, Rayvanth R. Chappidi, Bisiayo E. Fashemi, Paul A. Felder, Amy Mora, Sandra L. Grimm, Cristian Coarfa, Indira U. Mysorekar

**Author notes:** Lead contact: Indira U. Mysorekar, Ph.D., Department of Medicine, Section of Infectious Diseases, Baylor College of Medicine, One Baylor Plaza, 535E, Houston, Texas 77030. These authors contributed equally to this work (Chetanchandra S. Joshi, Arnold M. Salazar).

## Abstract

Aging is a risk factor for disease via increased susceptibility to infection, decreased ability to maintain homeostasis, inefficiency in combatting stress, and decreased regenerative capacity. Multiple diseases including urinary tract infection (UTI), are more prevalent with age; however, the mechanisms underlying how aging affects the urinary tract mucosa and the reason why aging correlates with disease are poorly understood. Here, we show that, relative to young (8-12 weeks) mice, the urothelium of aged (18-24 months) female mice accumulates large lysosomes with decreased acid phosphatase activity and shows overall decreased autophagic flux. Aged bladders exhibit basally high accumulation of reactive oxygen species (ROS) and dampened redox response. Furthermore, the aged urothelium exhibits a canonical senescence-associated secretory phenotype (SASP) at baseline with continuous NLRP3-inflammasome- and Gasdermin D (GSDMD)-dependent pyroptotic cell death. Accordingly, we find that aged mice chronically exfoliate epithelial cells. When infected with uropathogenic *E. coli*, infected aged mice harbor more bacterial reservoirs post-infection and are prone to spontaneous recurrent UTI. Finally, treatment of aged mice with D-Mannose, a natural bioactive monosaccharide, rescues autophagy flux, reverses SASP, and limits pyroptotic epithelial shedding. Thus, normal aging dramatically affects bladder physiology with aging alone increasing baseline cellular stress and susceptibility to infection. Additionally, our results suggest that mannose supplementation could serve as a senotherapeutic to limit age-associated urothelial dysfunction.

## INTRODUCTION

The average age of the world’s population is increasing. In 2018, the number of people aged 65 years or over outnumbered children under age five for the first time. By 2050, projections estimate there will be more than twice as many 65-year-olds than children less than 5 (United Nations et al., 2019). Aging impacts the organism at the cellular, tissue, and organ level, and is thus a risk factor for a plethora of chronic and severe diseases that arise mainly as a consequence of the organism’s decreased ability to maintain homeostasis or combat stress, increased susceptibility to infection (Mody & Juthani-Mehta, 2014; Rowe & Juthani-Mehta, 2013, 2014; Sadighi Akha, 2018), and decreased regenerative capacity (López-Otín et al., 2013). Aging also alters the immune system’s ability to mount both adaptive and innate immune responses, and promotes chronic stimulation of low-grade inflammation known as inflammaging (Ray and Yung, 2018). Overall, aging is marked by the accelerated loss of physiologic integrity, resulting in cellular dysfunction and increased susceptibility to death (López-Otín et al., 2013).

Aging also contributes significantly to increased risk for several diseases and conditions of the urinary tract, including urinary incontinence and urinary tract infections (UTIs), especially in postmenopausal women (Cotter et al., 2012; Dielubanza et al., 2014; Tsan et al., 2008). UTIs are the most common bacterial infection with more than 404.6 million cases reported worldwide in 2019 (Zeng et al., 2021). Recurrence is a major complication in UTI management, where a recurrent UTI (rUTI) is defined as at least three UTI episodes within 12 months or at least two episodes within 6 months (Kraft & Stamey, 1977). The main causative organisms of UTIs are uropathogenic *Escherichia coli* (UPEC) (Dielubanza & Schaeffer, 2011). To invade the urothelium, UPEC employ filamentous adhesive organelles called type 1 fimbriae with a terminal protein, FimH – a C-type lectin with high affinity for mannose residues on uroplakin receptors that line the urothelial cell surface (Mulvey et al., 1998; Ofek et al., 1977; Zhou et al., 2001). Following entry into fusiform vesicles from which they escape into the cytoplasm (Joshi et al., 2022; Pang et al., 2022), UPEC establish intracellular bacterial communities (IBCs) (Justice et al., 2012). Recognition of intracellular bacteria via host TLR-4/NLR receptors triggers the production of reactive oxygen species (ROS) (Joshi et al., 2021) and activates autophagic processes to expel bacteria into the bladder lumen (Terlizzi et al., 2017; Wang et al., 2012) thereby restricting further spread of UPEC. Urothelial cells can also be shed into the lumen to expel most of the UPEC-laden cells (Mulvey et al., 1998). Shedding further facilitates the recruitment of innate immune cells, including polymorphonuclear leukocytes, monocytes, and macrophages to attack the bacteria (Davis et al., 2005; Smith et al., 2008; Stemler et al., 2013). Immune infiltration, along with the inflammatory response, limits further invasion of UPEC. The bladder also activates antioxidant pathways via NRF2 to prevent high ROS build-up in the urothelium (Joshi et al., 2021; Wang et al., 2019), as this balance is essential to avoid excessive cell damage. However, despite the innate cellular and molecular responses employed by the bladder immune milieu, UPEC form quiescent intracellular reservoirs (QIRs), – bacterial communities that establish within terminally-differentiated superficial cells and/or underlying transitional epithelial cells by co-opting autophagy and persisting within autophagosomes for months and eventually re-start the cycle of infection (Wang et al., 2019; Wang et al., 2012; Wang et al., 2012).

Recent work has shown that aging alone initiates significant changes to the immune landscape of the aged female urinary tract including the formation of bladder tertiary lymphoid tissues (bTLT) (Hamade et al., 2022; Ligon et al., 2020). This raises important questions about how the urothelium is affected by aging and related immune changes, and what mechanistic role these changes might play in the development of UTIs and their recurrence. There are few previous studies of aging effects on key baseline aspects of the bladder’s ability to manage inflammation and respond to infection. In this study, using young and aged female mice with and without induced UTI, we provide evidence of significant cellular and molecular alterations in the aged bladder epithelium at baseline. Cytological and immunological assays and transcriptomic analyses revealed substantial alteration in regulation of autophagy, quantity of lysosomes, and mitochondrial function, concomitant to a dampened antioxidative response and an exacerbated immune response. Most importantly, correlating with what is observed in human patients, aged mice showed the effects of having increased number of bacterial reservoirs, including frequent spontaneous recurrence of infection, and increased urothelial shedding. We also show that the aged epithelium exhibits a senescence-associated secretory phenotype (SASP) and spontaneous, chronic pyroptotic cell death via activation of NLRP3 inflammasome and Gsdm D. Treatment of aged mice with D Mannose reverses the SASP and limits epithelial pyroptosis. Our data suggest the potential use of mannose supplementation as a senotherapeutic to treat age-associated urothelial dysfunction.

## RESULTS

### Aged bladders exhibit altered urothelial cellular architecture, and reduced autophagic flux and lysosomal function

Aged bladder epithelial cells showed accumulation of cytoplasmic vesicular structures (**Figure 1A**). Aged bladder tissue also showed, as expected, sub-epithelial follicular lymphocytic foci that we previously have characterized as age-induced tertiary lymphoid tissue (bTLT) (**Figure 1A**) (Ligon et al., 2020). In aged compared to young mouse urothelium, transmission electron microscopy (TEM) revealed a 4-fold increase in the number and 8-fold increase in mean size of lysosomes (**Figure 1B-D**). The accumulation of lysosomes in aged urothelium was underscored by staining with LysoTracker, a fluorescent dye that concentrates in lysosomes (**Figure 1E**). Similar lysosome congestion has been reported previously in aged rat bladder epithelia (Truschel et al., 2018). Lysosomal accumulation can indicate either increased lysosomal activity in the cell or decreased lysosomal hydrolase activity such that lysosomal function, and therefore, turnover is impaired. Thus, we probed lysosomal function using an assay to detect enzymatic activity of the lysosomal enzyme acid phosphatase (ACP). ACP activity was significantly lower in aged (∼64% reduction) compared to young bladders (**Figure 1F**), consistent with lysosomal congestion due to impaired function.

**Figure 1.**
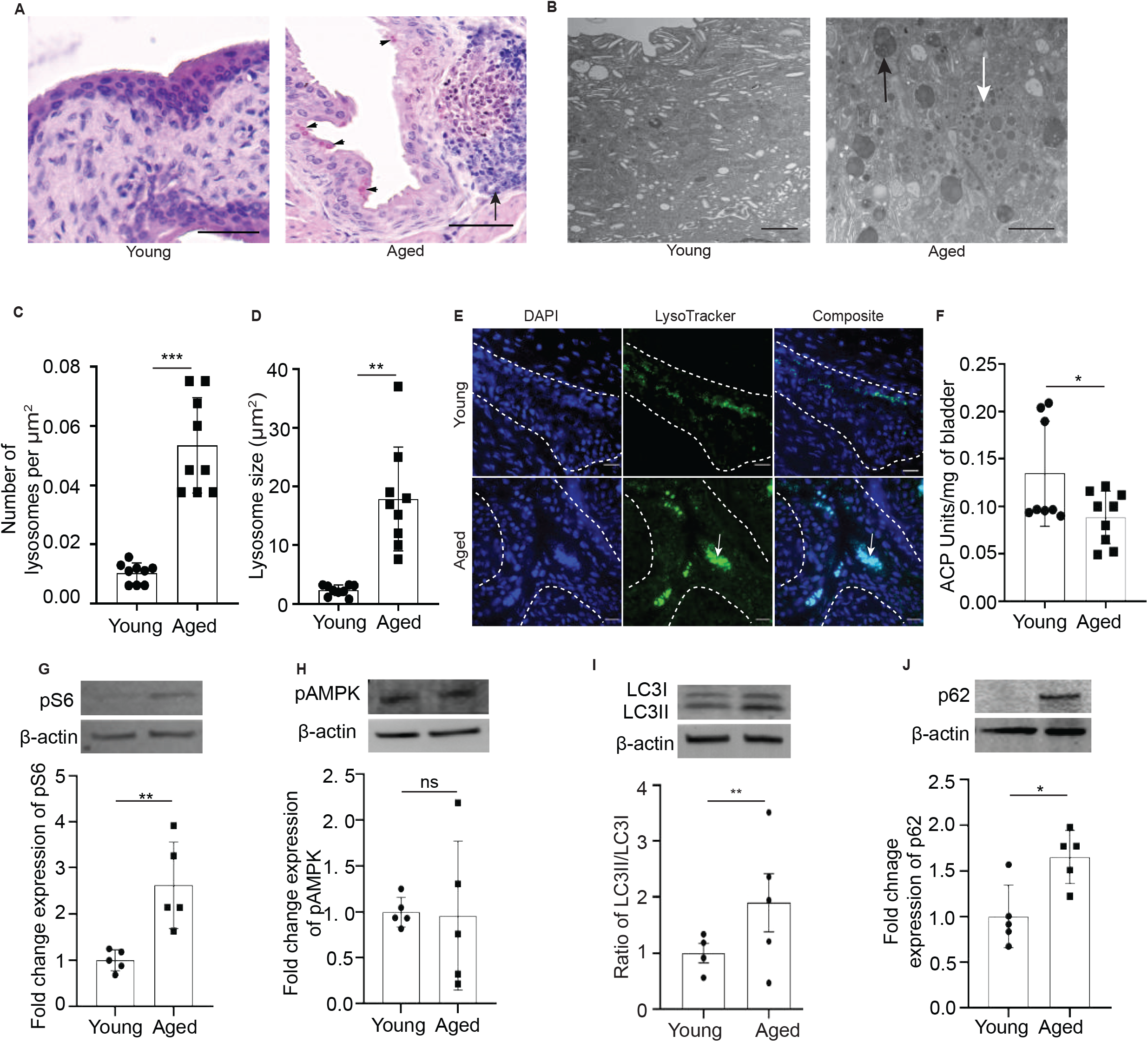
Aged bladders exhibit altered urothelial cellular architecture, and reduced autophagic flux and lysosomal function. **(A)** H&E staining of young and aged bladder. Arrowheads show sub-epithelial follicular foci, arrow indicates bTLT. Bar = 50 μm. **(B)** Transmission Electron Micrograph (TEM) of young and aged bladder. White arrow indicates lysosome, and the black arrow indicates enlarged lysosome. Bar = 2 μm. **(C)** Quantitation of the number of lysosomes per unit μm2 area. Data represented as mean ±SD (n = 9 images from total 5 animals in each group). ^***^p <0.001 compared with young urothelial sections by two-tailed unpaired t-test. **(D)** Quantitation of lysosome size in young and aged urothelium. Data represented as mean ±SD (n = 9 images from total 5 animals in each group). ^**^p <0.01 compared with young urothelial sections by two-tailed unpaired t-test. **(E)** Lysosome staining with LysoTracker (green) of young and aged urothelium. Arrows indicate the accumulated lysosomes in the aged urothelium. Bar = 20 μm. Nuclei are stained blue (DAPI). **(F)** ACP assay of indicated bladder tissue. Values are normalized to the bladder weight. Data are presented as mean ±SD (8 young and 9 aged bladders). ^*^p <0.05 compared with young by Mann-Whitney test. **(G)** Western blot and densitometric quantitation of pS6 in indicated tissue. Beta actin serves as a housekeeping control. Data are presented as mean ±SD (n = 5 animals in each group). ^**^p <0.01 compared with young by Mann-Whitney test. **(H)** Western blot and densitometric quantitation of pAMPK in indicated tissue. Beta actin serves as a housekeeping control. Data are presented as mean ±SD (n = 5 animals in each group). ns, non-significant compared with young by Mann-Whitney test. **(I)** Western blot and densitometric quantitation of LC3 I and II in young and aged bladder tissue. Beta actin serves as a housekeeping control. Data are presented as mean ± SD (n = 5 animals in each group). ns, non-significant, ^*^p < 0.05 compared with young by ordinary one-way ANOVA with Šídák’s multiple comparisons test. **(J)** Western blot and densitometric quantitation of p62 in indicated tissue. Beta actin serves as a housekeeping control. Data are presented as mean ±SD (n = 5 animals in each group). ^*^p <0.05 compared with young by Mann-Whitney test.

Lysosomal and mitochondrial responses and functions are tightly associated (Demers-Lamarche et al., 2016; Deus et al., 2020; Kim et al., 2021; Peng et al., 2020). Thus, we examined mitochondria in aged bladder cells. In almost all eukaryotes, mitochondrial range from 0.5 to 3μm, and their shape and number vary depending on cell types (Logan, 2010; Shami et al., 2021; Wiemerslage & Lee, 2016). MitoTracker staining revealed that the mitochondria in aged mouse urothelium were not only more abundant but were significantly larger (>8-fold) than those observed in a young bladder **(Figure S1)**. Lysosomal accumulation is often associated with dysregulated autophagy (Wang et al., 2019; Wang et al., 2012; Wang et al., 2012), which, in turn, is regulated by the mTORC1-AMP Kinase (AMPK) axis (Yang & Klionsky, 2010). The mTOR complex 1 (mTORC1) in particular suppresses autophagy (Wesierska-Gadek, 2010). Phosphorylation of the small ribosomal subunit 6 (phosphoS6) depends on mTORC1 phosphorylation of S6 kinase. Thus, phosphoS6 abundance in tissue is a faithful surrogate for mTORC1 activity (Willet et al., 2018). Aged bladders had ∼2.5-fold increase in phosphoS6 (**Figure 1G**) relative to young bladders, suggesting that aging was associated with increased mTORC1 activity. Activation of AMPK by phosphorylation was not consistently different between aged and young bladders (**Figure 1H**). To examine steady state autophagic flux, we measured the unlipidated (LC3I) and lipidated (LC3II) forms of LC3, the latter is a protein that is lipidated during autophagosome formation, and p62, a protein that targets cellular structures into autophagosomes. Aged bladders had increased LC3I and LC3II protein levels with a significant increase in LC3II/LC3I ratio (**Figure 1I**) and in p62 abundance (∼60%) (**Figure 1J**). LC3II requires active autophagy to form but requires completion of autophagy via lysosomal fusion and lysosomal enzymatic activity to be degraded, and p62 degradation also requires autophagosome-lysosome fusion and activity. Thus, the results are consistent with increased formation of autophagosomes and autolysosomes with decreased turnover of those structures and an overall pattern of increased mTORC1 activity with a block in late-stage autophagy.

### Aged urothelium exhibits senescence-associated secretory phenotype and a blunted antioxidant response

Next, we dissected the global status of transcripts in young and aged mice to evaluate the molecular signatures in the bladder in response to aging. RNA-Seq transcriptomic data revealed a robust and unique aged bladder signature (**Figure S2A**) with increased expression of 1,038 genes and decreased expression of 376 genes (**Figure S2B**). Gene set enrichment using the MSigDB Hallmark compendium revealed that in addition to changes in immune and inflammatory pathways, allograft cell/tissue rejection, cell death, oxidative stress, fatty acid synthesis, and mTOR/autophagy were identified as the top enriched pathways (**Figure S2C**). The elevation in immune inflammatory pathways was consistent with our previous report that described the increased tertiary lymphoid structures (Ligon et al., 2020).

Inflammation, oxidative stress, as well as increased lysosomal and mitochondrial size all occur during the cell-based program known as senescence (Correia-Melo et al., 2016; Fafián-Labora et al., 2019; Wiley et al., 2016). We reasoned that the aged urothelium could be undergoing a cellular senescence program even at steady state. Senescent cells display distinctive features like growth arrest, unresolved DNA damage, high senescence-associated β-galactosidase (SA-β-gal) activity, and pro-inflammatory SASP, which is a component of inflammaging (i.e., chronic inflammation that develops with advanced age, and can help spread senescence to adjacent cells) (Sabbatinelli et al., 2019). Consistent with senescence chronically occurring in aged bladder cells, we found increased SA-β-gal expression in aged compared to young bladders (**Figure 2A**). Senescent cells also shed portions of chromosomes from the nucleus, forming cytoplasmic chromatin fragments (CCFs) that can be marked by cytoplasmic accumulation of the DNA damage marker γ-H2AX (Hao et al., 2022; Ivanov et al., 2013). We observed that aged bladders exhibited high intensity expression of CCFs through γ-H2AX staining, further solidifying the presence of DNA damage response in aged bladder (**Figure 2B**). There was also increased expression of transcripts for the cell cycle arrest/DNA damage response genes p16 (∼80%), p21 (∼60%), and p53 (∼25%) in aged compared to young bladders (**Figure 2C-E**). Furthermore, multiplex analysis of urine samples from aged versus young mice showed a significant increase in cytokines, chemokines, and monocyte chemoattractant proteins, which recruit immune cells to clear senescent cells. We observed high expression of G-CSF, IL-1α, IL-6, IP-10, MCP-1, MIP-1α, and MIP-1β in aged compared to young bladders (**Figure 2F)**. Together, these findings demonstrate that the aged urothelium displays a senescence-associated secretory phenotype.

**Figure 2:**
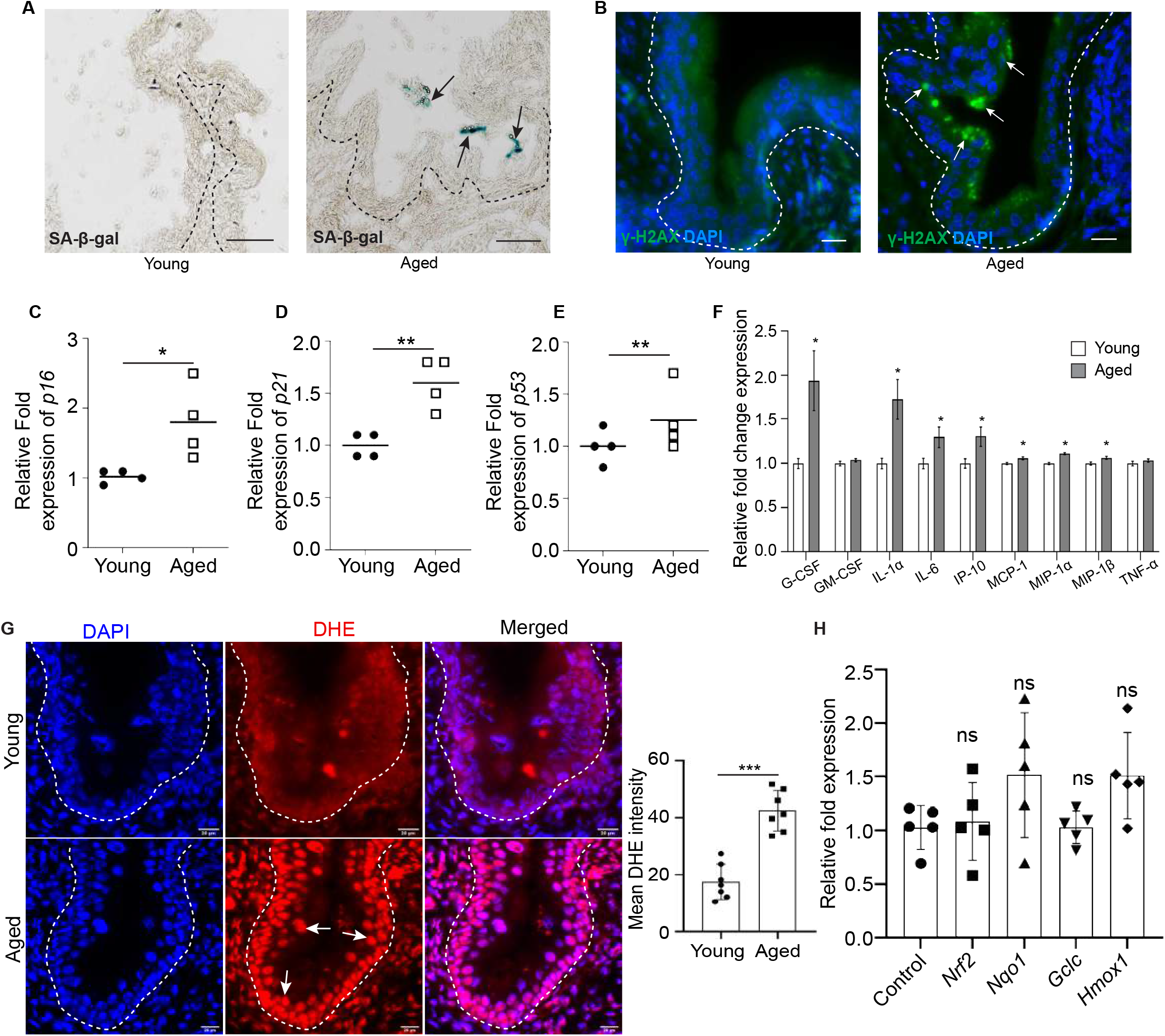
Aged urothelium exhibits senescence-associated secretory phenotype (SASP) and a blunted antioxidant response. **(A)** SA-β-gal staining in indicated bladder. Arrows indicate blue staining in aged bladder. Bar = 100 μm. **(B)** γ-H2AX localization (green) in indicated bladder, arrows indicate staining in aged bladder. Bar = 20 μm. Nuclei are stained blue (DAPI). **(C-E)** qRT-PCR analysis of p16, p21, and p53 in young and aged mice bladder. Data are presented as mean ±SD (n = 4 animals in each group). ^*^p <0.05, ^**^p <0.01 compared with young bladder by Mann-Whitney test. **(F)** Milliplex assay indicating fold change in the expression of SASP cytokines in young and aged urine. Data are presented as mean ±SD (n = 5 animals in each group). ^*^p <0.05 compared with young urine by Mann-Whitney test. **(G)** ROS indicator dye Dihydroethidium (DHE) (red) staining in young and aged bladder. Arrows indicate nuclear DHE. Nuclei are stained blue (DAPI). Bar = 20 μm. Quantitation of DHE stained area in young and aged bladder. Data represented as mean ±SD (n = 7 images from total 7 animals in each group). ^***^p <0.001 compared with young urothelial sections by two-tailed unpaired t-test. **(H)** qRT-PCR analysis of Nrf2 and its targets Nqo1, Gclc, and Hmox1. Data are presented as mean ±SD (n = 5 animals in each group). ns non-significant compared with control (Young bladder) by two-tailed unpaired t-test.

We previously showed that altered autophagy or lysosomal function leads to impaired antioxidant response resulting in the accumulation of ROS in the context of UTIs (Joshi et al., 2021). To investigate the effect of aging on ROS production, we used dihydroethidium (DHE), a dye that hydrolyzes upon interaction with ROS and stains the nucleus. This revealed high signal intensity in the aged urothelium, and quantification of the signal indicated higher levels of ROS (∼100% higher) in the aged urothelium relative to the low baseline levels in the young bladders (**Figure 2G**). However, despite markedly elevated ROS, the normal cellular antioxidant response genes (*Nrf2, Nqo1, Gclc*, and *Hmox1*) were not increased relative to control levels (**Figure 2H**), indicating ROS may continuously accumulate in aged bladders because of diminished antioxidant response. Overall, these findings demonstrate that aged bladder epithelial cells constitutively induce a cellular senescence program even in an otherwise unperturbed state.

### Aging is associated with increased NLRP3/Gsdm/IL-1β axis in the urothelium and pyroptotic epithelial cell death

Next, we sought to further characterize the mechanisms contributing to the hyper-inflammatory state and consequences of a senescent and oxidatively stressed bladder. The NLRP3 inflammasome is an important component of the innate immune system that modulates caspase-1 activation and the secretion of pro-inflammatory cytokines in response to microbial infection in epithelial cells and macrophages (Demirel et al., 2020; Kelley et al., 2019; Orning et al., 2019; Symington et al., 2015; Wu et al., 2019). Together with our whole bladder RNA Seq data showing multiple gene pathways involved in inflammatory responses and cell death, we reasoned that excessive ROS and SASP could trigger a baseline inflammatory and cell damage response. Indeed, we observed that aged bladders have significantly elevated levels of *Nlrp3* (**Figure 3A**). Given that NLRP3 as part of the inflammasome that converts inactive pro-caspase 1 into its mature form, we investigated the levels of pro- and cleaved caspase-1 in young and aged bladders and found significant accumulation of the mature form in aged bladders (**Figure 3B**). Caspase-1 cleaves pro-IL-1β into IL-1β (Denes et al., 2012; Sollberger et al., 2014; Vijayaraj et al., 2021). Accordingly, we found that pro-IL-1β and cleaved IL-1β were both significantly higher in aged compared with young bladders (**Figure 3C**). Caspase-1 cleaves a family of proteins called Gasdermins, which play a pivotal role in releasing IL-1β by forming membrane pores that increase cell membrane permeability and induce pyroptosis (Orning et al., 2019). We detected increased expression of transcripts for *GsdmA, GsdmC, GsdmD*, and *GsdmE* (**Figure 3D**). The Human Protein Atlas indicates that the predominant Gasdermin protein expressed in human bladder epithelium is GsdmD (Tissue Expression of GSDMD - Summary - The Human Protein Atlas). Thus, we further investigated protein expression of GsdmD in aged murine bladders and found significantly increased expression of cleaved GsdmD resulting in N-terminal (GsdmD-NT) fragments (**Figure 3E**) that form the membrane pore through, where IL-1β is released. Thus, the aged bladder exhibits high levels of the NLRP3 inflammasome and a steady-state caspase-1/IL-1β/GsdmD cascade (**Figure 3F**).

**Figure 3:**
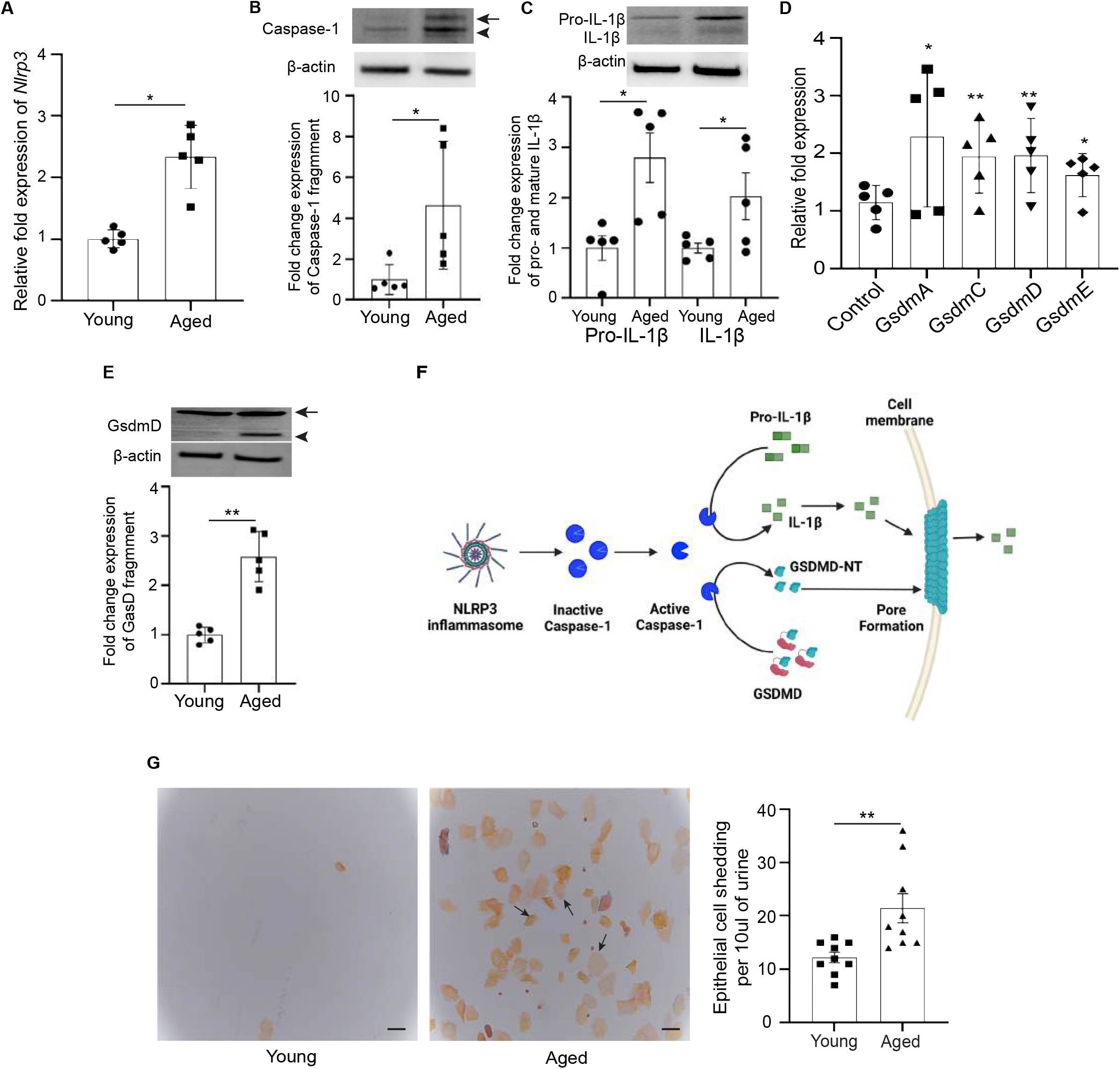
Aging is associated with increased NLRP3/Gsdm/IL-1β axis in the urothelium and pyroptotic epithelial cell death. **(A)** qRT-PCR analysis of Nlrp3. Data are presented as mean ±SD (n = 5 animals in each group). ^*^p <0.05 compared with young control bladder by Mann-Whitney test. **(B)** Western blot and densitometric analysis of Caspase-1. Arrow indicates full-length protein; arrowhead shows cleaved fragment. Densitometric analysis was done for the cleaved fragment. Beta actin serves as a housekeeping control. Data are presented as mean ±SD (n = 5 animals in each group). ^*^p <0.05 compared with young by Mann-Whitney test. **(C)** Western blot and densitometric analysis of pro-IL-1β and IL-1β. Beta actin serves as a housekeeping control. Data are presented as mean ±SD (n = 5 animals in each group). ^*^p <0.05 compared with young by Mann-Whitney test. **(D)** qRT-PCR analysis of GsdmA, GsdmC, GsdmD, GsdmE. Data are presented as mean ±SD (n = 5 animals in each group). ^*^p <0.05, ^**^p <0.01 compared with young control bladder by Mann-Whitney test. **(E)** Western blot and densitometric analysis of GasderminD. Arrow indicates full-length protein, arrowhead shows N-terminal cleaved fragment. Densitometric analysis was done for the cleaved fragment. Beta actin serves as a housekeeping control. Data are presented as mean ±SD (n = 5 animals in each group). ^**^p <0.01 compared with young by Mann-Whitney test. **(F)** Graphical representation of GasderminD-mediated pore formation and pyroptosis. **(G)** Representative urine cytology images from uninfected young, aged and D-mannose treated aged mice. Arrows show exfoliated bladder epithelial cells. Bar = 100 μm. Quantitation of exfoliated/shed epithelial cells in young and aged mice. Data represented as mean ±SD (n = 9 animals in each group). ^**^p <0.01 compared as indicated by two-way ANOVA Bonferroni’s multiple comparisons test.

Next, we determined if the constitutively increased NLRP3/GsdmD/IL-1β axis in the urothelium would affect tissue homeostasis. Indeed, cytological examination of urines from young and aged bladders revealed a significant level of spontaneously shed epithelial cells in aged urines (∼80% increase) (**Figure 3G**). This finding is similar to what has been demonstrated in urine samples from aged/post-menopausal women (Meister et al., 2021). Thus, together our results suggest that in the aged bladder, high levels of NLRP3 inflammasome activates a caspase-1/IL-1B/GsdmD cascade leading to pyroptotic cell death resulting in constitutive epithelial shedding.

### Aging exacerbates UTI recurrence

Postmenopausal/elderly women have a higher risk and frequency of UTIs than younger pre-menopausal women. We next evaluated whether the epithelial and immune changes we have delineated in our aged bladders would impact UTI susceptibility and/or pathogenesis. Young and aged mice were infected with a clinical UTI isolate, UPEC strain UTI89, and the impact on infection was determined. Initial urine titers between young and aged mice were similar (**Figure 4A**). However, infected aged bladders were robustly inflamed compared with infected young bladders 24 hours post infection (hpi) (**Figure 4B**) with increased influx of mononuclear cells observed in urine (∼100% increase) (**Figure 4C**). We then sought to determine how aging-associated inflammatory response and cellular changes affects the pathogenesis of UPEC in the urothelium by counting the intracellular bacterial communities (IBCs) in infected mice (Justice et al., 2012). Compared to young mice, infected aged mice showed a 50% increase occurrence of IBCs, detected as IBC-containing superficial cells shed in urine during the time 6 to 72h time window of an acute UTI (**Figure 4D-E**).

**Figure 4.**
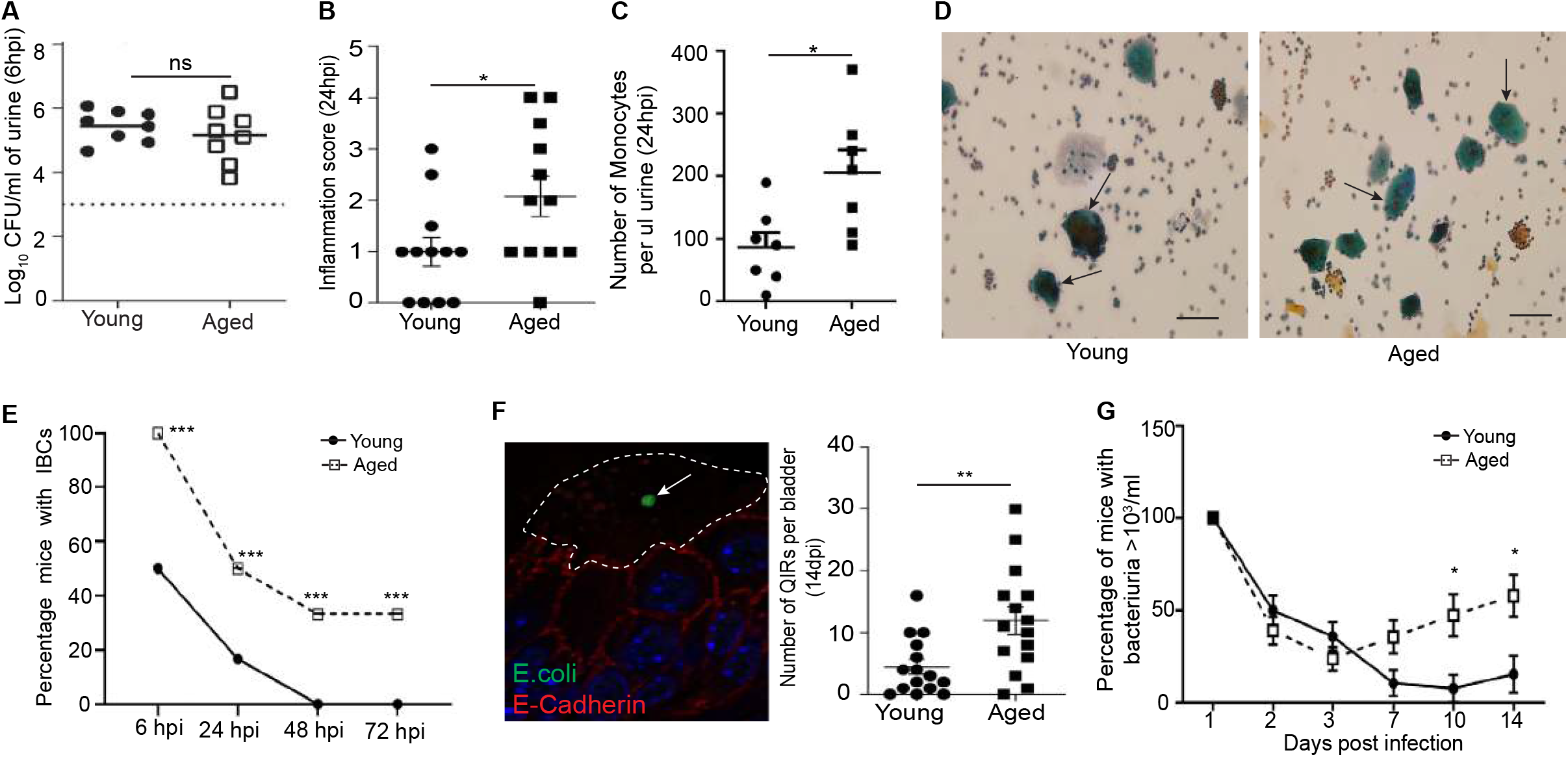
Aging exacerbates UTI recurrence. **(A)** Quantitation of bacterial load in indicated mice urine at 6 hpi. Data are presented as mean ±SD (n = 8 mice in each group). ns non-significant compared with the young by two-tailed unpaired t-test. **(B)** Inflammation score of urines obtained from indicated mice post-infection. Data are presented as mean ±SD (n = 12 animals in each group). ^*^p <0.05 compared with young by two-tailed unpaired t-test. **(C)** Quantitation of the number of monocytes in urine in indicated mice post infection. Data are presented as mean ±SD (n = 7 animals in each group). ^*^p <0.05 compared with young by two-tailed unpaired t-test. **(D)** Representative urine cytology images of young and aged mice at 24 hpi. Arrows indicate infiltrated immune cells; arrowheads indicate cells loaded with IBC. Bar = 50 μm. **(E)** Quantitation of the number of IBC formed per bladder in indicated mice at indicated time post infection. Data are presented as mean ±SD (n = 6 at 6 hpi, n = 12 at 24 hpi, n = 6 at 48 hpi, n = 6 at 72 hpi in young and aged mice). ^***^p <0.001 compared with young mice bladder by two-way ANOVA Bonferroni’s multiple comparisons test. **(F)** Quantitation of the number of QIRs formed per bladder in indicated mice at 14 days post infection (dpi). Data are presented as mean ±SD (n = 15 animals in group). ^**^p <0.01 compared with young mice bladder by two-tailed unpaired t-test. Immunofluorescence localization of *E. coli* (green) forming QIR (arrow) in aged bladder. E-cadherin (red) marks cell membrane. Dotted area encircles superficial cell. Bar = 10 μm. **(G)** Quantitation of percentage of young and aged mice with bacteriuria at indicated times post-infection. Data are presented as mean ±SD. For young group, n is 1 dpi = 38, 2 dpi = 38, 3 dpi = 39, 7 dpi = 19, 10 dpi = 13, 14 dpi = 13, for aged group n is 1 dpi = 41, 2 dpi = 41, 3 dpi = 42, 7 dpi = 28, 10dpi = 19, 14dpi = 19. ^*^p <0.05 compared with young mice by two-way ANOVA Bonferroni’s multiple comparisons test.

We have previously shown that a subset of UPEC can persist within epithelial cells in the intact epithelium in the form of QIRs (Mysorekar & Hultgren, 2006). Quantification of QIRs at 14 days post infection (dpi) in infected mice using an *E. coli*-specific antibody revealed a significant increase in the number of QIRs in infected aged bladder relative to infected young bladders (**Figure 4F**). Finally, commensurate with increased QIRs that can seed recurrent infections, a significantly higher percentage of infected aged mice exhibited spontaneous recurrence of bacteriuria (**Figure 4G**). Together, our findings demonstrate that the uninfected aged bladder is significantly more inflamed. Aged bladders also develop more IBCs suggesting increased bacterial invasion and intracellular proliferation. Though IBCs are shed, QIRs remain within the epithelium to seed recurrent infection, explaining why aged mice are susceptible to spontaneously recurrent UTI.

### D-mannose treatment ameliorates age associated SASP, inflammation, and epithelial pyroptosis

Several clinical investigations have described the benefits of D-mannose in patients suffering from rUTI (Chiu et al., 2022; Kranjčec et al., 2014; Parazzini et al., 2022). D-mannose is known to bind to UPEC adhesin protein FimH and thus prevent them from adhering to bladder epithelial cells (reviewed in Lupo et al., 2021). However, D-mannose has also been shown to dampen inflammation (Ito et al., 2022). Here, we sought to test the hypothesis that D-mannose decreases inflammation in aged bladders. We treated aged mice with 1.1M D-mannose in drinking water for 14 days. As expected, D-mannose administration did not have any adverse effects on weight or overall health of the mice (data not shown). A subset of aged mice who were infected with UPEC for 14 days and treated with D-mannose alone also did not show a change in urinary bacterial load, consistent with D-mannose not having bacteriostatic or bactericidal activity (data not shown). However, in contrast to the significantly increased LC3II/LC3I ratio in aged bladders compared with young bladders, D-mannose-treated aged bladders showed a significant reduction in the LC3II/LC3I ratio when compared with untreated aged mice (**Figure 5A**). A significant reduction in p62 (∼65%) was also observed in the D-mannose-treated aged mouse bladders (**Figure 5A**), indicative of re-activated autophagic flux. Furthermore, we observed a significant decrease in the level of β-galactosidase (∼55%) (**Figure 5B**) suggesting reduced SASP. Accordingly, we observed reduced transcript levels of p21 (∼45%) and p53 (∼65%) in the D-mannose-treated aged bladder compared with the untreated group although p16 levels remained unchanged (**Figure 5C**). Most notably, however, we observed a significant decrease in spontaneous epithelial cell shedding from uninfected D-mannose-treated aged mice urine compared with uninfected non-treated aged mice (∼75% decrease) (**Figure 5D-E**). Finally, cytological quantification of inflammatory cells in urine samples revealed a lowered inflammation score in D-mannose-treated aged mice when compared with the untreated aged group (**Figure 5F**). Our findings suggest that D-mannose could be considered as a potential senotherapeutic approach as it can significantly improve age-associated dysfunction by rescuing the block in autophagic flux, SASP, inflammation, and urothelial cell pyroptosis. Our study adds promise to the utility of D-mannose that goes beyond its use in treating urinary tract infections alone.

**Figure 5.**
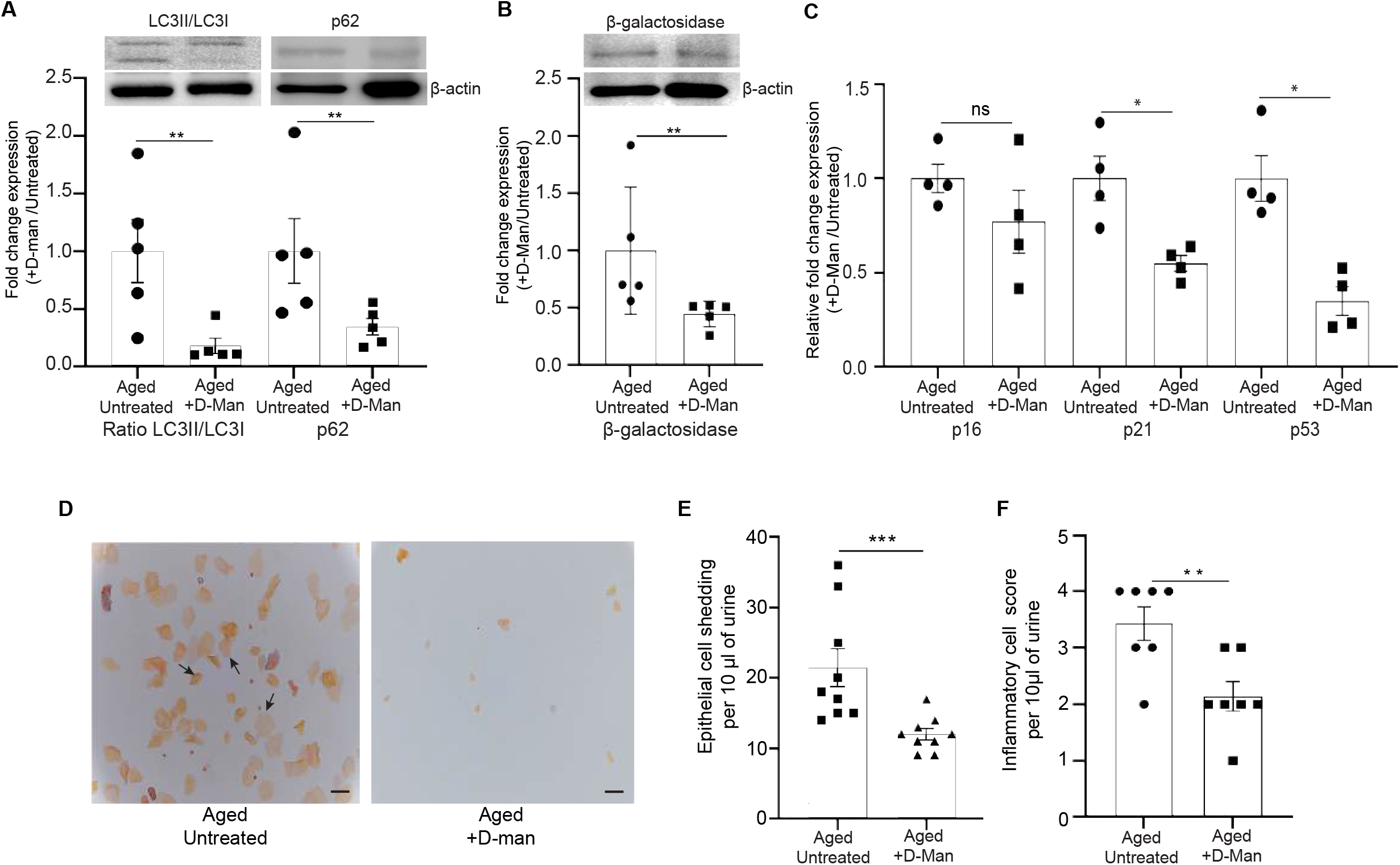
D-mannose treatment ameliorates age-associated SASP, inflammation, and epithelial pyroptosis. **(A)** Western blot and densitometric analysis of LC3, and p62. Beta actin serves as a housekeeping control. Data are presented as mean ±SD (n = 5 animals in each group). ^*^p <0.05, ^**^p<0.01 compared with untreated aged bladder by Mann-Whitney test. **(B)** Western blot and densitometric analysis of Beta-galactosi-dase. Beta actin serves as a housekeeping control. Data are presented as mean ±SD (n = 5 animals in each group). ^**^p <0.01 compared with untreated aged bladder by Mann-Whitney test. **(C)** qRT-PCR analysis of p16, p21, and p53 in D-mannose treated and untreated aged bladders. Data are presented as mean ±SD (n = 4 animals in each group). ns non-significant, ^*^p <0.05 compared with untreated aged bladder by Mann-Whitney test. **(D)** Representative image of urines from untreated and D-mannose-treated aged mice, Bar = 100 μm. **(E)** Quantitation of exfoliated/shed epithelial cells in untreated and D-mannose-treated aged mice. Data represented as mean ±SD (n = 9 animals in each group). ^***^p <0.001 compared as indicated by two-way ANOVA Bonferroni’s multiple comparisons test. **(F)** Inflammatory cell score of urines obtained from aged D-mannose-treated and untreated mice. Data are presented as mean ±SD (n = 7 animals in each group). ^**^p <0.01 compared with aged by Mann-Whitney test.

## DISCUSSION

The aging population is steadily increasing, creating increasing need to provide healthcare and treatments capable of meeting the increased susceptibility to infectious diseases and other conditions that occurs in aging. In the lower urinary tract, especially in women, aging combined with postmenopausal status exacerbates multiple conditions of the urinary tract including recurrent urinary tract infections (rUTIs). There is a great need to understand impact of aging and age-related changes in the urinary tract mucosa, and to delineate the potential mechanisms driving this increased susceptibility. At the tissue and cell level, the aging process is associated with altered intercellular communication and dysfunction, loss of protein homeostasis, and cellular senescence among a myriad of other alterations (López-Otín et al., 2013; Sadighi Akha, 2018). Cellular senescence has been directly implicated as a key driver of aging and aging-related diseases. However, little is known on how these processes occur and/or affect disease outcomes in the bladder mucosa. In this study, we demonstrate that bladder urothelial cells undergo significant cellular and biochemical modifications simply because of normal aging alone, including the development of SASP, sustained oxidative stress, lysosome dysfunction, and decreased autophagic flux. We show that aging is also associated with increased inflammation, pyroptotic cell death, and susceptibility to rUTIs. We further demonstrate that oral administration of D-mannose, a natural bioactive monosaccharide, limits SASP and inflammatory death of urothelial cells.

Lysosomes are cellular organelles that contain enzymes to break down waste materials and cellular debris in post-mitotic cells. Over time, lysosomes progressively accumulate oxidatively modified macromolecules and defective organelles that are not fully degraded. Lysosomal dysfunction is commonly observed in age-related neurodegenerative diseases including Alzheimer’s and Parkinson’s and impaired lysosomal activity has been shown to play an important role in the development of these disorders (Koh et al., 2019; Navarro-Romero et al., 2020). We found significant modifications in the lysosomal component of the uninfected aged urothelium, including low acid phosphatase activity, and increased number and size of lysosomes. This indicates a compensatory response that enhances the digestion of accumulating cellular debris/waste, ameliorating inefficiency or abnormality in lysosomal functions. Our findings align with previously reported work showing accumulation of an aging pigment, lipofuscin, in aged urothelial cells (Perše et al., 2013), decreased acidification, loss of cathepsin B activity, and expanded endolysosomal compartments in the aged rat urothelium (Truschel et al., 2018).

A decrease in lysosomal degradation can lead to altered autophagy, including defective mitophagy, further leading to the accumulation of ROS and disrupted cellular function. Indeed, we noted an increase in the size and number of mitochondria in addition to lysosomes in the aged urothelium. Cellular respiration and mitochondrial bioenergetics in aged urothelial cell cultures have recently been shown to have a reduced mitochondrial Ψm, oxygen consumption rate, and ATP release (de Rijk et al., 2022). Mitochondrial accumulation during high oxidative conditions is also attributed to autophagy dysfunction (Luo et al., 2013).

We and others have shown that dysfunctional autophagy leads to downregulation of the antioxidant response (Joshi et al., 2021; Qiao et al., 2020; Yun et al., 2020; Zhao et al., 2019). In agreement with previous findings (de Rijk et al., 2022), we report here that the aged bladder has elevated levels of ROS. If mitochondrial accumulation leads to more ROS and more oxidative damage, one could expect that the antioxidative process would be activated to counteract high oxidative stress (Joshi et al., 2021). However, the levels of antioxidant response-related genes were not activated in aged mice compared to young mice, suggesting an ineffective or a defective antioxidant response, which may lead to an unregulated increase of inflammation in aged bladders. Higher ROS activity that exacerbates oxidative damage could lead to apoptosis or senescence (Bladier et al., 1997; Chen et al., 1998). Thus, aging of the bladder overall increases the initiation or activation of a constitutively stressed phenotype.

In this study, we further determined that the aged urothelium displays multiple hallmarks of cellular senescence including increased SA-β-galactosidase activity, and the acquisition of a pro-inflammatory SASP. Senescent cells can exhibit a compromised nuclear envelope allowing chromatin fragments to bud off and enter the cytoplasm to be degraded via an autophagosome-lysosome process (Ivanov et al., 2013). Lysosomes play an important role in limiting generation of SASP, which is in part driven by the presence of such cytoplasmic chromatin fragments (CCF) (Lee et al., 2006; Miller et al., 2021; Young et al., 2009). Reduced autophagic flux and lysosomal function that was evident in our aged bladder epithelial cells would suggest that chromatin fragments could accumulate in the cytoplasm. Indeed, we observed the presence of γ-H2AX positive CCF in cytoplasm of aged urothelial cells but not in young cells, suggesting activation of the DNA stress pathway and presence of CCF in senescent urothelial cells.

Accumulation of DNA in the cytoplasm serves as a potent danger signal that can activate an innate immunity cytosolic DNA sensing pathway, leading to proinflammatory responses. In fact, SASP is associated with various growth-regulatory factors, cytokines, and chemokines (Krtolica et al., 2001; Mylonas & O’Loghlen, 2022). We found a significant increase in secreted soluble factors such as IL-1α, IL-6, and MIP-1α in aged mouse urines and tissues. The role of cytoplasmic DNA in age-associated inflammation in several diseases is beginning to be described. For example, premature aging syndrome such as ataxia telangiectasia (AT), a severe neurodegenerative syndrome, is characterized by accumulation of γ-H2AX-enriched cytoplasmic DNA along with the expression of inflammatory genes, including IL-6 (Lan et al., 2019; Miller et al., 2021; Song et al., 2019). Previous work from our group has shown that the aged urothelium has increased permeability characterized by exposure of underlying urothelial cells and the stromal compartment to urinary content (Sawhill et al., 2022), which is highly detrimental to the normal urothelial function of maintaining an impermeable barrier to waste products and cytotoxic factors in urines (Dalghi et al., 2020). We posit that chronic exposure to urinary cytotoxic irritants could trigger DNA damage secondary to accumulation of ROS and drive CCF formation, leading to an SASP-associated inflammatory response coupled with age-associated dysfunctional lysosomal function and mitophagy defects.

The outcomes of these age-associated changes appear to be inflammatory/lytic cell death of the urothelial cells via activation of an inflammasome/Gasdermin/IL-1β axis triggered by the NLRP3 inflammasome. Activation of this lytic cell death pathway (pyroptosis) has been shown to lead to pro-inflammatory cytokine IL-1β release (Yu et al., 2021). This occurs via assembly of gasdermins (Gsdm), a family of cell death effectors with highly conserved N and C terminal domains and a variable linker region that oligomerize to form pores in the cell membrane, and induce cell permeabilization (Broz et al., 2020). In our study, we determined that all known Gsdm orthologs in mice (GsdmA, GsdmC, GsdmC, GsdmD, and GsdmE) showed significant upregulation of transcript levels in the uninfected aged bladders. The most prominent member of the family, GsdmD and specifically the active fragment GsdmD-NT, also had high protein expression levels in the aged bladder. To our knowledge, this is the first report on Gasdermin expression in the urothelium and its upregulation with age and role in spontaneous pyroptotic cell death. We and others have previously shown that UPEC infection induces IL-1β mediated pyroptosis in bladders and in macrophages (Symington et al., 2015; Wu et al., 2019). Our study suggests that aging itself, as a condition, is sufficient to activate pyroptosis and facilitate the release of pro-inflammatory cytokines by increased activation of NLRP3, caspase-1, and IL-1β akin to an acute infection condition. Pyroptosis has been implicated in the etiology of nerve cell injury and death in neurodegenerative diseases including Alzheimer’s disease and Parkinson’s disease (Yue et al., 2022). The changes we have noted in the aging urothelium could constitute urothelial degeneration and underlie a number of bladder diseases. We suggest that mechanisms of senescence and pyroptosis may be common to multiple tissues with age and should perhaps be investigated in a pan-tissue manner.

The excessive shedding of urothelial cells captured in urines of aged female mice in the current study and noted previously in postmenopausal women (Meister et al., 2021) could be responsible in part for the dysregulated urothelial barrier function (Sawhill et al., 2022). The regenerative capacity of the urothelium in response to injury/infection/stress is notoriously robust (Murray et al., 2021; C. Wang et al., 2017; Wiessner et al., 2022). However, it is likely that the aged urothelium may be compromised in its capacity to regenerate new luminal umbrella cells to replenish the continuously shedding cells into the urine. Our findings that aged urothelial cells present with increased cell cycle arrest/DNA damage response genes (p16 and p21) supports the notion that urothelial cells undergo senescence and are likely unable to fully restore barrier damage. Interestingly, activated p21 has been shown to further induce ROS production, thus reinforcing the response. Given our findings that the antioxidant response via Nrf2 and its cognate pathway components is compromised, the constitutive stress and shedding cycles could significantly impact barrier function. Indeed, Nrf2 signaling is known to protect cells from pyroptosis by attenuating the activation of the NLRP3 inflammasome (Kensler et al., 2007). Furthermore, we have demonstrated that *Nrf2*^-/-^ mice also exhibit spontaneous urothelial shedding in urine and the shed cells are highly ROS-positive (Joshi et al., 2021). Thus, urothelial shedding in response to unmitigated ROS could represent an inability to repair damaged/stressed cells. This aging phenotype coupled with the inability to restore the lost cells could underlie multiple urological disease processes. Detailed understanding of urothelial regenerative potential in the aged bladder is warranted to develop interventions to block ROS, limit SASP, or promote rejuvenation of the urothelium.

Frequency of recurrent UTIs is significantly higher in postmenopausal women (Al-Badr & Al-Shaikh, 2013; Jung & Brubaker, 2019; Raz, 2011; Zhu et al., 2020). There are multiple hypotheses for the increase including reduction of systemic estrogen levels, reduced detrusor activity, and an altered urobiome, among others (Alperin et al., 2022). Recent work from our group and others has identified a new and potentially critical contributor to an altered bladder mucosal environment with age that is responsible for inflammaging, namely, formation of bladder tertiary lymphoid tissue (bTLT) with bone fide germinal centers that arise in an age-dependent manner coincident with loss of fertility in female mice (Ligon et al., 2020). Notably, bTLT only forms in a sex-dependent manner in female bladders (Hamade et al., 2022). bTLT occurs in postmenopausal women and is significantly associated with increased frequency of rUTIs and shorter time intervals between recurrent episodes (Ligon et al., 2020) and are associated with bacteria including *E. coli* (De Nisco et al., 2019). We note an increased incidence of rUTIs in our aged mouse bladders, including formation of more cytoplasmic bacterial biofilm communities as determined by IBC-containing umbrella cells shed in the urine. Interestingly, we have also noted a significantly higher number of QIRs which form within autophagosomes (Mysorekar & Hultgren, 2006) and can spontaneously emerge to give rise to a new infection cycle, resulting in spontaneous bacteriuria as observed in post-menopausal women (Meister et al., 2021). We saw an increased LC3I/II ratio with age and a concomitant block in autophagy flux, which suggests that there is accumulation of autophagosomes that could provide a greater number of protected niches for QIR formation. Our findings that aged bladders infected with UPEC exhibit a spontaneous rUTI phenotype that is not seen in young infected C57Bl6/J mice is consistent with our clinical findings (Ligon et al., 2020). The mechanistic and direct links between our murine studies and clinical findings of bTLT and increased frequency of rUTIs need to be better understood. Moreover, the links between urothelial dysfunction with age and inflammaging and cellular senescence remain to be solidified. Our bulk whole bladder transcriptomics data indicating increases in inflammatory response and cell death pathways and decreases in pathways associated with protein degradation and epithelial mesenchymal transition might provide further clues to understanding interdependent urothelial-immune crosstalk and implications for disease pathogenesis.

Finally, we report on the utility of a simple sugar monosaccharide, D-mannose, as a surprisingly effective senotherapeutic in limiting pyroptotic epithelial cell death, reversing cell cycle arrest, limiting SASP, and restoring autophagic flux that is otherwise notably increased in non-treated aged mice. D-Mannose is a sugar that is considered as an alternative treatment to UTIs (Chiu et al., 2022; Lenger et al., 2020) as it binds to the adhesion protein FimH on uropathogens and prevents their adherence to bladder epithelial cells (Scribano et al., 2020) and has been known to decrease secretion of inflammatory cytokines (Ala-Jaakkola et al., 2022). Our results show that D-mannose has a cell intrinsic impact in addition to the known extrinsic roles, as treatment reduces accumulation of SA-β-galactosidase and activates autophagic flux. Interestingly, treatment also affects cell cycle regulators, p21and p53 although not p16. p21 is mainly activated early in the initiation of senescence, whereas p16 maintains cellular senescence (Huang et al., 2022). Thus, D-mannose might block early steps in the process of senescence and thereby limit further progression. This would make D-mannose a highly effective seno-morphic as it modifies senescence risk. In line with this, mannose has been reported to protect against intestinal barrier dysfunction in a (young) mouse model of colitis (Dong et al., 2022). Moreover, we have recently reported that D-mannose prophylaxis led to a significant decrease in UTI incidence rate in patients with bTLT (Chiu et al., 2022). Thus, D-mannose treatment might produce far-reaching urothelial cell intrinsic and extrinsic reversal of age-associated changes in addition to limiting rUTI frequency. As the decrease in shedding, cell death, and senescence point to rehabilitation or improvement in the overall cellular and molecular status of the aging bladder, further investigation is warranted to understand the ways D-mannose treatment contributes to these processes, and the potential benefits of D-mannose beyond rUTI prevention and treatment.

## METHODS

### Mice

C57B6/J female mice (8-12 weeks: young; and 18-24 months: aged) were obtained from the National Institute of Aging and maintained at Washington University School of Medicine and Baylor College of Medicine. Standard rodent chow and water were available *ad libitum* throughout the experiment. Mice were housed in groups of five in a temperature-(22 ± 1 °C) and humidity-controlled vivarium with lights maintained on a 12:12 light/dark cycle. All animal experimental procedures were approved by the Institutional Animal Care and Use Committee at Washington University School of Medicine (Animal Welfare Assurance #A-3381-01) and Baylor College of Medicine (Animal protocol number AN-8629). All mice were humanely euthanized at the end of each experiment.

### UPEC strain and infection

A clinical cystitis isolate of uropathogenic *E. coli*, UTI89, was grown in Luria-Bertani (LB) broth overnight in static culture at 37 °C incubator the day before infecting mice. Adult female mice (2 months to 18 months of age) were anesthetized and transurethrally-inoculated with 10^7^ colony forming units (CFU) of UTI89 suspended in PBS as previously described (Hung et al., 2009).

### D-Mannose treatment

Aged female mice were treated with drinking water supplement with 1.1M D-mannose for 14 days (Zhang et al., 2017). Control cohort was provided with un-supplemented water. Weights of the mice were recorded, and urine samples were collected for cytological preparation and analysis. Water consumption was also recorded/calculated.

### H&E staining

Bladder sections from young and aged mice were deparaffinized by soaking in 3 separate solutions of 100% Histoclear at 5 minutes each. Sections were rehydrated in decreasing concentrations of ethanol (100%, 90%, 70%, 50%) at 5 minutes per concentration. Staining of the nuclei was performed by soaking the sections in Hematoxylin solution for 5 minutes followed by rinsing in running tap water to remove excess Hematoxylin solution. Sections were dipped quickly into an acid alcohol solution, soaked for 5 minutes in sodium bicarbonate solution, followed by rinsing in distilled water. Staining of the cytosol was done immediately by dipping the slides 3 times in Eosin solution, soaking in increasing concentrations of ethanol (50%, 70%, 90%, 100%) at 3 minutes each, followed by a final dip in Histoclear. Sections were applied with xylene-based mounting medium, covered with cover glass, and edges were sealed with clear nail polish. Image capture was performed with a Panoramic Midi microscope (3DHISTECH Ltd, Hungary).

### Transmission Electron Microscopy (TEM)

Young and aged mouse whole bladders were processed as described previously (Wang et al., 2012) to examine lysosomes and lipid droplets. The bladders were fixed with fixative containing 2% glutaraldehyde and 3% PFA in 0.1M sodium cacodylate. Samples were rinsed three times in sodium cacodylate buffer and post-fixed in 1% osmium tetroxide for 1 hour, stained in 1% uranyl acetate for 1 hour, rinsed and dehydrated, and subjected to critical point drying. Samples were then gold-coated and analyzed on a JEOL-1200 EX II Transmission Electron Microscope (JEOL, USA). For count and size determination of lysosomes, 9 images of five mice in each group were analyzed using ImageJ analysis software.

### Lysosome Staining

Fixed, fresh-frozen sections of young and aged bladders were utilized for staining lysosomes with Lysotracker Red (L7528, Life Technologies) following manufacturer’s protocol with modifications. Sections were soaked and incubated in 100nM Lysotracker Red probe solution for 30 mins at 37 °C, rinsed in 1X PBS, and counterstained with DAPI. Sections were applied with Xylene-based mounting medium, covered with cover glass, and sealed edges with clear nail polish. Images were captured using Zeiss Axio Imager M2 Plus Wide Field Fluorescence Microscope (Carl Zeiss Inc, Thornwood, NY).

### Mitochondria Staining

Fixed, fresh-frozen sections of young and aged bladders were utilized for staining of mitochondria with Mitotracker dye (M22426, Thermo Fisher Scientific, USA) following manufacturer’s protocol with modifications. Sections were soaked and incubated in 100 nM Mitotracker dye solution for 5 minutes at 37°C, rinsed in 1X PBS, and counterstained with DAPI. Sections were applied with Prolong Gold Antifade mountant (P10144, Thermo Fisher Scientific, USA), covered with cover glass, and edges were sealed with clear nail polish. Images were captured using Zeiss Axio Imager M2 Plus Wide Field Fluorescence Microscope (Carl Zeiss Inc, Thornwood, NY). Quantitation of mitochondrial size was performed on 5 images from 5 mice in each group using ImageJ.

### Oil Red O Staining

Frozen sections of young and aged mice bladder were fixed in 10% neutral buffered formalin for 15 minutes, rinsed with distilled water, and air-dried. Sections were dehydrated in 60% isopropyl alcohol, stained in Oil Red O solution for 20 minutes at room temperature, dipped in 60% isopropyl alcohol, and rinsed in distilled water. Sections were counterstained with hematoxylin and mounted in glycerol jelly. Images were captured using Zeiss Axio Imager M2 Plus Wide Field Fluorescence Microscope (Carl Zeiss Inc, Thornwood, NY). Lipid droplets were quantitated from 4 sections of 4 mice from each group using ImageJ.

### RNA-Seq Analysis

Tissue preparation and RNA sequencing protocols were adapted from Ligon et al (2020). Briefly, snap-frozen mice bladders were homogenized to isolate RNA using RNeasy Mini Kit (74101, Qiagen) and RNase-free DNase digestion kit (79254, Qiagen) following manufacturer’s protocol. Libraries were prepared with Ribo-Zero rRNA depletion kit (Illumina) and sequenced on a HiSeq3000 (Illumina). Reads were aligned to the Ensembl GRCm38.76 top-level assembly with STAR version 2.0.4b and gene counts were derived from the number of uniquely aligned unambiguous reads by Subread: featureCount version 1.4.5. Sequencing performance was then assessed for total number of aligned reads, total number of uniquely aligned reads, genes detected, ribosomal fraction, known junction saturation, and read distribution over known gene models with RSeQC version 2.3. Gene expression expressed as counts was normalized using upper quartile normalization and RUVr (Risso et al., 2014), then differentially expressed genes were determined using the EdgeR R package (Robinson et al., 2010). Significance was achieved for FDR-adjusted p-value<0.05 and fold change exceeding 1.5x. Enriched pathways were determined using over-representation analysis (ORA) as implemented by the MSigDB online platform (Liberzon et al., 2014), specifically using the hypergeometric distribution against the MSigDB Hallmark pathway collection (Liberzon et al., 2015), with significance achieved for FDR-adjusted p-value<0.05. Data was visualized using the R statistical system and GraphPad Prism.

### Quantitative RT-PCR (qRT-PCR)

Bladders of 5 young and 5 aged mice were collected and homogenized in TRIzol™ Reagent (15596026, Thermo Fisher Scientific, USA) to extract total RNA followed by DNase 1 treatment (18068-015, Thermo Fisher Scientific, USA) following manufacturer’s protocol. One microgram of total RNA was utilized to perform cDNA synthesis using SuperScript™ II Reverse Transcriptase (18064-014, Thermo Fisher Scientific, USA) following manufacturer’s protocol. All cDNAs were diluted to 1:8 with RNase-free water prior thereafter. Primer designs (see Key Resources Table) and qRT-PCR setup was performed using SsoAdvanced Universal SYBR® Green Supermix (1725274, Bio-Rad, USA) following manufacturer’s protocol in 10 μl reactions (5 μl Supermix; 1 μl each of Forward and Reverse Primers; 2 μl diluted cDNA; 1 μl RNase-free water), each reaction was done in triplicate. 18s rRNA was used as a housekeeping gene. The qRT-PCR reaction was run in QuantStudio™3 Real-Time PCR System (Applied Biosystems™, USA) using the following settings: 98 °C, 3 minutes (initial activation); 98 °C, 30 seconds (Denaturation); 58 °C, 30 seconds (Annealing/Extension); 40 cycles; and instrument default setting for melt-curve analysis. Raw quantitation values were used for calculating fold change based on the young bladder.

### Milliplex (Cytokine Assay)

SASP-specific cytokines (G-CSF, GM-CSF, IL-1α, IL-6, IP-10, MCP-1, MIP-1α, MIP-1 β, and TNF-α) were quantitated using the Milliplex kit (MCYTOMAG-70K, MilliporeSigma, Germany). Second void urine samples from 5 animals in each group were collected and processed the same day in duplicate following the manufacturer’s protocol. Microplate reading was performed using BIOPLEX 200 (Biorad, USA) at Baylor College of Medicine Antibody-Based Proteomics Core Laboratory. Intra-assay %CVs (coefficient of variation) were calculated for GM-CSF (8.03%), IL-1α (9.13%), IL-1β (7.17%), IL-6 (8.10%), MIP-1α (9.79%), and TNF-α (6.96%). In addition, creatinine assay (MAK080, MilliporeSigma, USA) was performed on the same urine samples (3μl) following the manufacturer’s protocol. Cytokine concentrations were normalized to the corresponding average creatinine level of young and aged mice.

### SA-β-galactosidase Staining

Fresh-frozen 8 uM sections of young and aged bladders were utilized for SA-β-galactosidase staining. Briefly, sections were fixed in 1% formamide solution for 5 minutes at room temperature and washed twice in 1X PBS for 3 minutes each. Staining was done using a solution composed of the following: 40 mM citric acid/sodium phosphate buffer (pH 6.0), 150 mM NaCl, 2 mM MgCl2, 5 mM potassium hexacyano-ferrate II, 5 mM potassium hexacyano-ferrate III, and fresh 1 mg/mL 5-bromo-4-chloro-3-indolyl-β-D-galactosidase (X-gal) dissolved in dimethylformamide (DMF) at 37 °C (without CO2) overnight. Image capture was performed with a Panoramic Midi microscope (3DHISTECH Ltd, Hungary).

### γ-H2AX Staining

Formalin-fixed paraffin-embedded (FFPE) bladder sections from young and aged mice were deparaffinized by soaking in 3 separate solutions of 100% Histoclear at 5 minutes each. Sections were rehydrated in decreasing concentrations of ethanol (100%, 90%, 70%, 50%) at 5 minutes per concentration and soaked in 1X PBS for 5 minutes. Sections were blocked in 1% bovine serum albumin for I hour at room temperature, followed by antibody staining with anti-γH2AX (1:500; 9718, Cell Signaling Technology) in blocking buffer plus 0.1% Tween-20 overnight at 4°C. Sections were rinsed in 1X PBS (3 times, 5 minutes each), and incubated in secondary antibody AF488 (A11034, Thermo Fisher Scientific, USA) for 1 hour at room temperature and rinsed in 1X PBS (3 times, 5 minutes each). Sections were applied with Prolong Gold Antifade reagent with DAPI (P36935, Thermo Fisher Scientific, USA), covered with cover glass, and edges were sealed with clear nail polish. Sections were imaged under ECLIPSE Ni Epi-fluorescence Upright Microscope (Nikon, USA).

### Western Blotting

Bladders of 5 young and 5 aged mice were collected and homogenized in RIPA lysis buffer to extract total proteins. Total protein was quantitated using a BCA assay (Pierce™ BCA Protein Assay Kit, 23225, Thermo Fisher Scientific, USA). 10 ug of protein was loaded onto precast gel (4561095, 4–20% Mini-PROTEAN® TGX™ Precast Protein Gels, Bio-Rad, USA) and resolved at 200 volts for 30-35 minutes. Proteins bands were immobilized through transfer to PVDF membrane (IPFL00010, Immobilon Transfer Membrane, Millipore, Ireland) for one hour at 110 volts in ice. The membrane was treated with blocking buffer (927-60001, Intercept® (TBS) Blocking Buffer, LI-COR, USA) for 1 hour at room temperature with gentle agitation. Membranes were incubated with primary antibody solution in blocking buffer plus 0.1% Tween-20, overnight at 4°C with agitation. Beta-actin was used as a loading control. Membranes were washed in 1X TBS with 0.1% Tween-20, 5 times, for 5 minutes each at room temperature with agitation followed by treatment with appropriate secondary antibody solution in blocking buffer plus 0.1% Tween-20 for 1 hour at room temperature with agitation. Membranes were washed in 1X TBS with 0.1% Tween-20, 5 times, for 5 minutes each at room temperature with agitation followed by washing in 1x TBS 2 times, for 5 minutes each. Imaging was performed using the ChemiDocTM MP imAging system (Bio-Rad, USA). Densitometric analysis was done using Bio-Rad Image Lab software (6.0.1).

### Acid Phosphatase Assay

Fresh-frozen sections of young and aged bladders were tested for acid phosphatase activity using an acid phosphatase assay kit (CS0740-1KT, Sigma-Aldrich) following the manufacturer’s protocol. The experiment was carried out in 8 young and 9 aged bladders. Values were normalized to the bladder weight.

### *In vivo* ROS Assay and Imaging

OCT-embedded fresh young and aged bladders were cut into 10 μm thickness and mounted in glass slides for staining. Sections were soaked in ROS-sensing dye (Dihydro-Ethidium: DHE), incubated for 10 minutes at room temperature, washed in 1X PBS, and followed by counterstaining with DAPI. Images were captured using Zeiss Axio Imager M2 Plus Wide Field Fluorescence Microscope (Carl Zeiss Inc, Thornwood, NY). ROS signal in 7 images from 7 mice per groups was quantitated using ImageJ.

### Urine analysis and Inflammation Score

Young and aged mouse urine samples were collected for sediment analysis (10 μl urine plus 40 μl 1X PBS) and subjected to cytospin3. Sediments on the microscope slides were fixed in acetic acid/alcohol for 15 minutes and subjected to Epredia™ Papanicolaou EA Staining (22-050-211, Thermo Fisher Scientific, USA) following the manufacturer’s protocol. Stained urine sediments were examined using blind scoring under the light microscope. An inflammation score scaled from 0 to 4 was adopted, where 0 indicated <1 and 4 indicated >20 polymorphonuclear leukocytes per high-powered field (as previously described in Stemler et al., 2013). Exfoliated cells were also quantitated and expressed per 10 μl of urine, including the number of monocytes at specified hpi.

Urinary bacterial load from infected bladders of young and aged mice were quantified as described previously (Wang et al., 2012a). Briefly, urine samples were collected at indicated time points and serially diluted in 1X PBS. LB plates were spotted with 5 μL of each dilution 6 times, and plates were incubated at 37 °C overnight. Bacterial titers were calculated as log_10_ CFU/ml (6 hpi). Calculation of mice with bacteriuria was also performed.

### Pap Staining of IBCs in Urine

Urine from infected bladders of young and aged mice was fixed using acetic/alcohol fixative and dehydrated in 95% alcohol, followed by rehydration in water. Hematoxylin was incubated for 10 minutes and developed in running tap water. Sections were dehydrated in 95% alcohol and stained with brand OG-6 solution (from 22-050-211, Thermo Fisher Scientific, USA) for 2 minutes and washed with 95% alcohol. Next, EA-65 stain (from 22-050-211, Thermo Fisher Scientific, USA) was used for 10 minutes, followed by a dehydration step. Clearing was done in xylene and a xylene-based mounting media was used to mount the sides, followed by imaging under a Panoramic Midi microscope (3DHISTECH Ltd, Hungary).

### Quantification of QIRs

Quantification of QIRs was done as described previously (Wang et al., 2012). Briefly, young and aged mice were sacrificed at 14 days post infection (dpi) and bladders were collected and processed. Eight separate 5 μm step sections over a thickness of 300 μm were stained with antibodies against *E. coli*, and E-cadherin. Sections were imaged at 63S oil on a Zeiss Apotome microscope and processed using ImageJ. The total number of UPEC reservoirs were counted and reported per bladder (n=15 animals in each group).

### Statistical Analyses

All measured values were plotted using GraphPad Prism version 9.0.1 (GraphPad Software, La Jolla, CA, USA). Data were expressed as mean ± Standard Deviation (SD) of the sample size (n) (indicated in each figure). Unpaired Student’s t-tests, Mann-Whitney and ANOVA were used to determine statistical significance.

## Acknowledgments

This work was supported in part by NIH grants, R01DK100644, R01AG052494, P20DK119840, and R56AG064634 (to IUM), NIH training grants, T32AI007172 and T32GM007200 (to MML); and T32-AI007172 (to BEF). SLG and CC are partially supported by the Cancer Prevention Institute of Texas (CPRIT) RP170005, RP200504, and RP210227, NIH/NIAID 1U19AI144297, NIH/NCI P30 shared resource grant CA125123, and NIEHS grants P30 ES030285 and P42 ES027725. We thank Dr. Wandy Beatty at Molecular Microbiology Imaging Facility, Washington University School of Medicine for expertise with TEM and Drs. Jason Mills and Robert Lawrence for valuable comments and editing.

## Author contributions

CSJ, AMS, and IUM conceived the experimental plan; CSJ, AMS, CW performed the majority of experiments and were assisted by RRC, BEF, MML, PAF, AM. SLG and CC analyzed the RNA-Seq data. CSJ, AMS, and IUM wrote the manuscript, and all authors approved the final draft.

## SUPPLEMENTARY FIGURE LEGENDS

**Figure S1:**
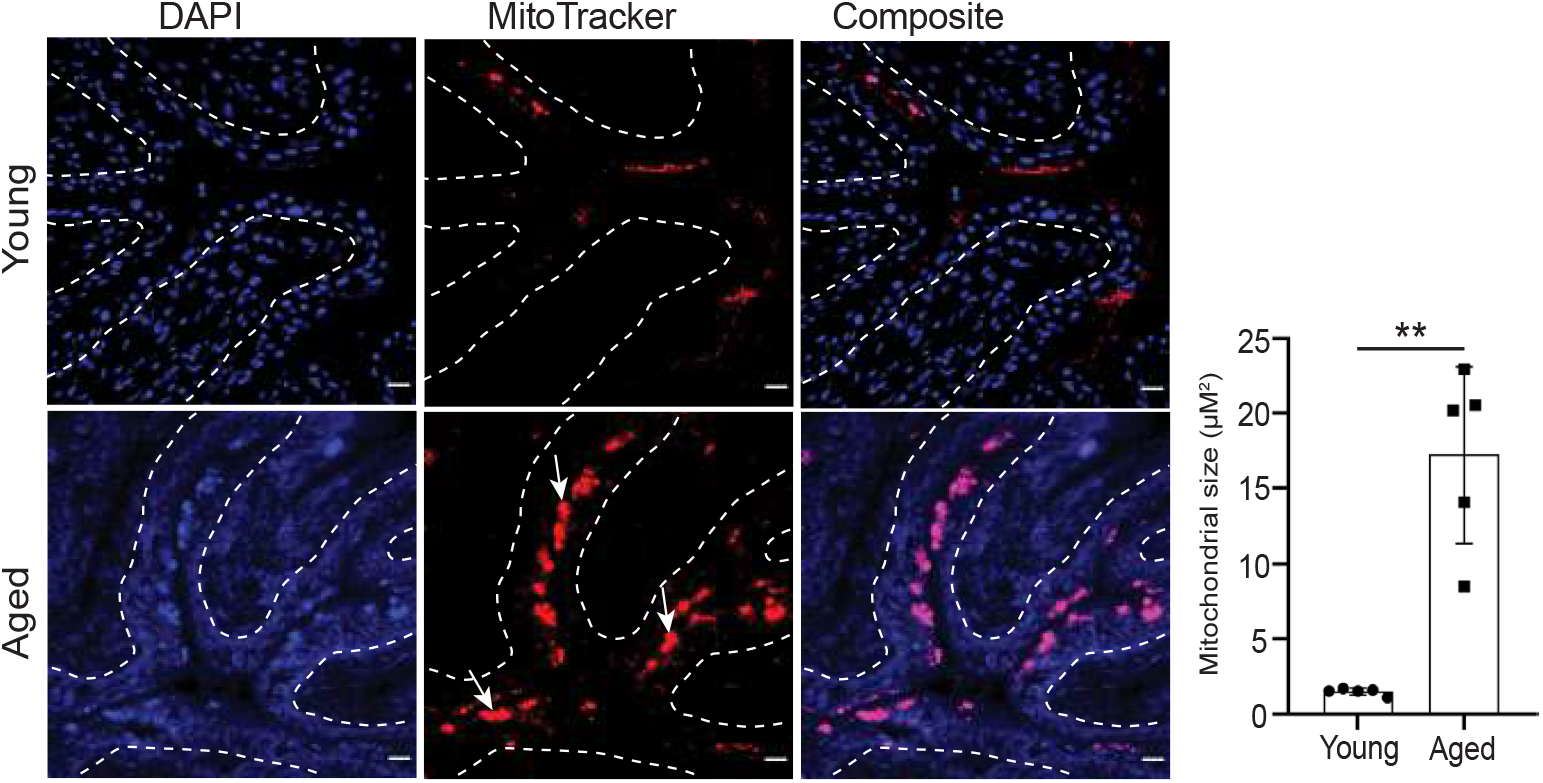
Mitochondrial staining and quantitation of mitochondrial size using MitoTracker (red) of young and aged urothelium. Arrows indicate the enlarged mitochondria in the aged urothelium. Bar = 20 μm. Nuclei are stained blue (DAPI). Graph data represented as mean ±SD (n = 5 images from total 5 animals in each group). ^**^p <0.01 compared with young urothelial sections by two-tailed unpaired t-test.

**Figure S2:**
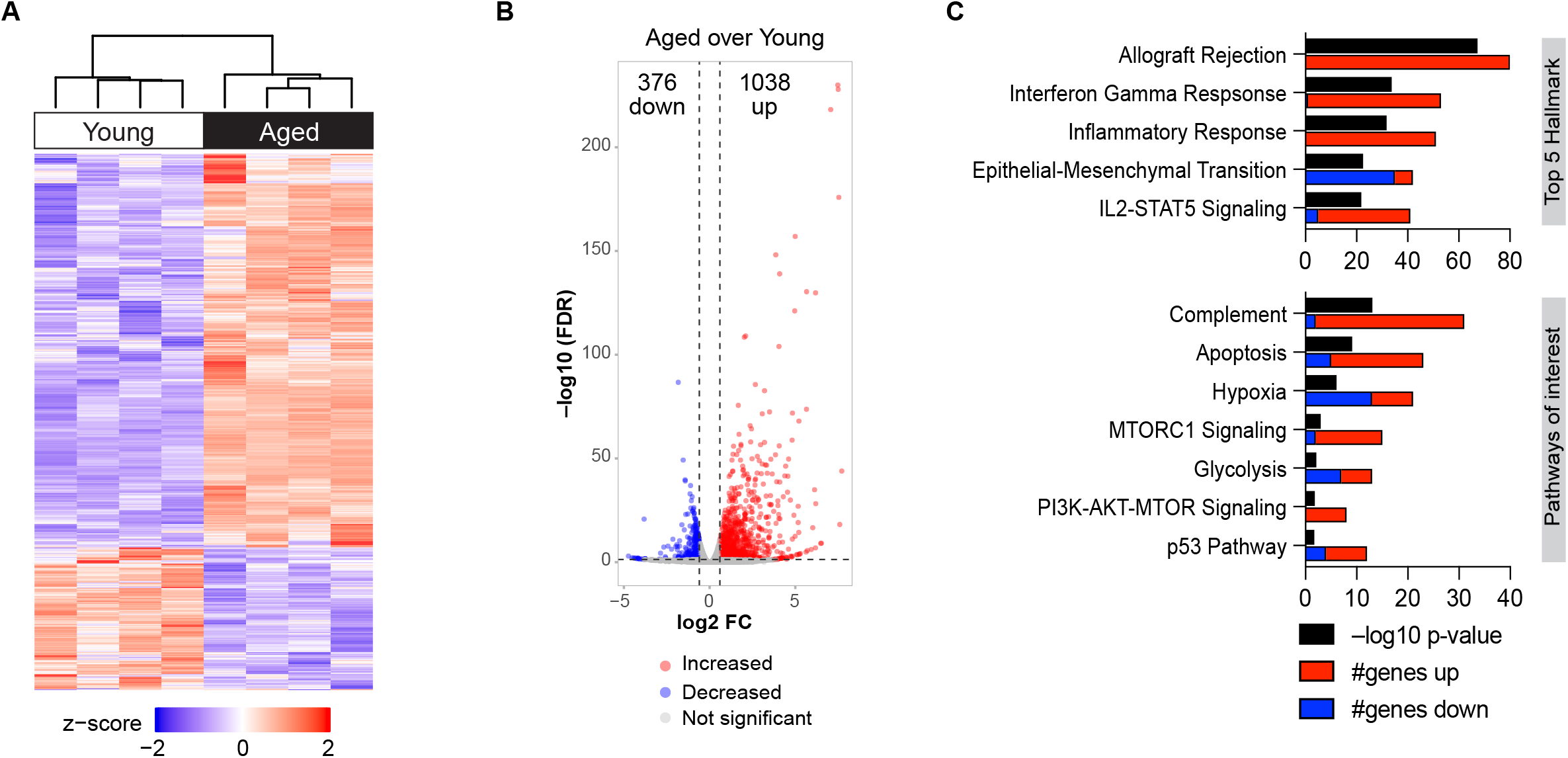
Global transcriptomic analysis of young and aged urothelium. **(A)** Heatmap of the global tissue transcriptome of young and aged bladders with fold change exceeding 1.5 and Benjamini-Hochberg FDR-adjusted p <0.05. n = 4 bladders/group. **(B)** Volcano plot of differentially expressed genes in aged compared to young bladder **(C)** Significantly enriched Hallmark pathways based on fold change expression of related genes.

